# ChexMix: A Literature Content Extraction Tool for Bioentities

**DOI:** 10.1101/2021.03.09.434525

**Authors:** Heejung Yang, Beomjun Park, Jinyoung Park, Jiho Lee, Hyeon Seok Jang, Namgil Lee, Hojin Yoo

## Abstract

Biomedical databases grow by more than a thousand new publications every day. The large volume of biomedical literature that is being published at an unprecedented rate hinders the discovery of relevant knowledge from keywords of interest to gather new insights and form hypotheses. A text-mining tool, PubTator, helps to automatically annotate bioentities, such as species, chemicals, genes, and diseases, from PubMed abstracts and full-text articles. However, the manual re-organization and analysis of bioentities is a non-trivial and highly time-consuming task. ChexMix was designed to extract the unique identifiers of bioentities from query results. Herein, ChexMix was used to construct a taxonomic tree with allied species among Korean native plants and to extract the medical subject headings unique identifier of the bioentities, which co-occurred with the keywords in the same literature. ChexMix discovered the allied species related to a keyword of interest and experimentally proved its usefulness for multi-species analysis.

## Introduction

The currently enhanced computing power is boosting the acquisition and processing of scientific data obtained from wet and dry lab experiments. In the fields of biology and chemistry, a huge amount of literature is being published every day and uploaded to the databases in real time. Several databases are currently available applying manual curation or *in silico* approaches for data management in arbitrary forms^1–5^. Hence, finding the meaningful information from large databases is almost like ‘finding a needle in a haystack’. Therefore, automation techniques, such as text-mining and natural language processing (NLP) methods, were developed to help convert raw scientific texts into well-structured scientific data^6,7^. PubTator Central (PTC) is a state-of-the-art text-mining service for automated annotation of bioentities including genes/proteins, genetic variants, diseases, chemicals, species, and cell lines in about 30 million abstracts and 3 million full-text articles available in PubMed (https://www.ncbi.nlm.nih.gov/pubmed)^8,9^. The bioentities co-occurring in the body of literature and extracted by PTC can be re-organized as a complex network and used for literature-based discovery.

Herein, we introduce ChexMix, a bioentity extraction tool for the extractions of interrelationships between medical subject headings (MeSH, https://www.ncbi.nlm.nih.gov/mesh) terms and taxonomy identifiers (TaxIDs) in the National Center for Biotechnology Information (NCBI) Taxonomy^10^. ChexMix is an open source python module for accessing and processing various forms of data from multiple biomedical databases. It collects the bioentities, such as species, chemicals, and diseases, which were annotated by the PTC, from abstracts and/or free full-text biomedical articles stored in the PubMed database. ChexMix converts and links the bioentities found in the literature with the unique identifiers of the species (TaxID), chemicals (MeSH), and diseases (MeSH). The association between these bioentities can be modulated into bipartite or multipartite networks, or hierarchical trees, aiding to inspect and simply understand the holistic structures of the information associated with targeted topics queried as keywords.

Each bioentity has its own hierarchical organization system according to the bioentity type. For taxonomy, species is located in the lowest rank of the taxonomic hierarchy and is involved with the higher ranks, genus and family. The hierarchical location and distance in the taxonomic tree between the species bioentities provide clues to discover other hidden bioentities. Species under the same genus share more similar features compared with different species under another genus. For chemicals, since similar structures can interact with proteins holding comparable binding pockets, the types of backbones or derivatives help inspect their physicochemical properties and/or biological roles. Moreover, annotation methods are introduced to classify the structural types of chemicals in chemicals profiling studies^11–13^. Even in the case of protein targets, the expression of genes helps understand the physiological function of proteins, as well as to identify related physiological disorders and pathologies^14,15^. ChexMix was developed to extract the bioentities based on the literature keywords, as well as the keywords entered by researchers (Fig. 1). Therefore, ChexMix was designed to help organize biomedical data into hierarchical knowledge based on topological similarities between bioentities. The co-occurrence of biomedical terms provides the assumption that the bioentities in the same abstract or full-text can be considered to be biologically or chemically related to each other^2^. These associations can then be visualized as network graphs or hierarchical trees, and be more easily analyzed for uncovering hidden insights from already existing knowledge.

**Figure 1.**
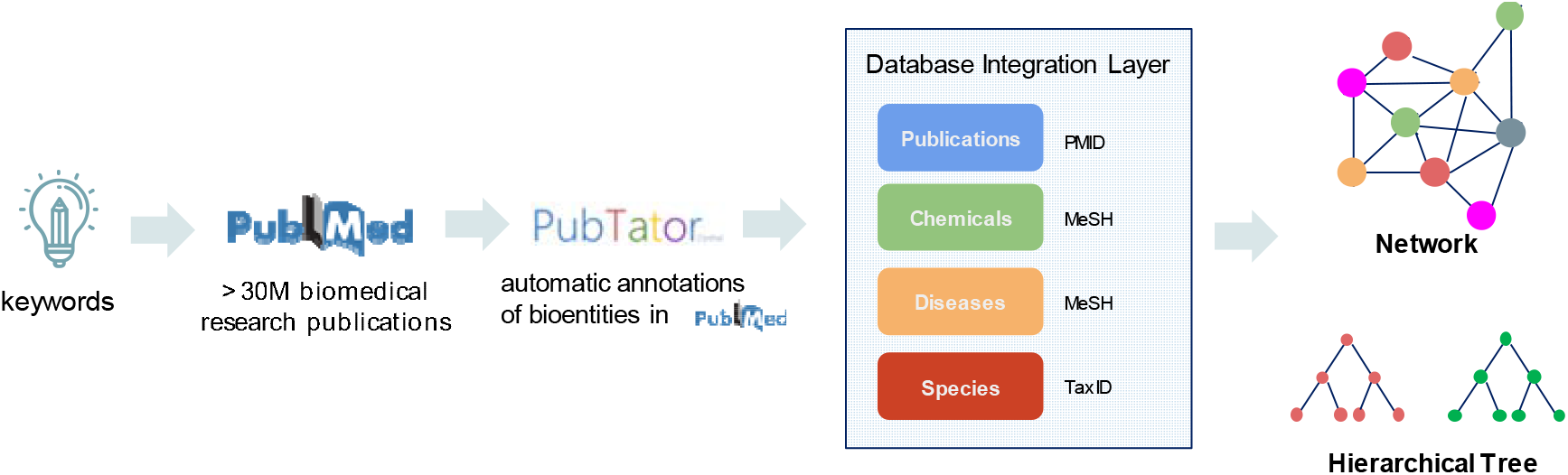
Network and hierarchical tree of biomedicals using ChexMix.

## Results and Discussion

ChexMix was designed for the extraction of hierarchical and topological information related to bioentities. Therefore, ChexMix extracts the bioentities that co-occur with the keywords queried in PubMed and encodes into unique identifiers indexing their related information. The combination of a hierarchical representation with a mapping of bioentities to identifiers at each level allows the relationships between them to be organized and cross-referenced. For example, species resulting from keywords of interest, such as chemicals or diseases, can be hierarchically represented from the highest rank, ‘cellular organisms’, according to the phylogenetic taxonomic system of the NCBI taxonomy database^10^. The search results are arranged according to the hierarchical characteristics of each bioentities and can be displayed in plots for hierarchical data visualization or nested lists (Fig. 1); therefore, the information can be useful for the inspection of related information among keywords of interest. Herein, ChexMix was applied to discover the biomedical sources of natural products that produce the bioactive compound, amentoflavone, which holds a wide range of biological activities, including antioxidative, anti-inflammatory, anticancer, antiviral, and antifungal properties^17^. This compound also shows potent antisenescence activity against ultraviolet B irradiation-induced skin aging, preventing nuclear aberrations^18^; thus, it can be used for the prevention of skin aging in the cosmetic industry.

Firstly, 319 bioentities were extracted from ChexMix using the keyword ‘amentoflavone’ under the highest taxonomic rank, ‘cellular organisms’ (Fig. 2). Among them, 223 species comprised in the Viridiplantae (literally ‘green plants’) clade were targeted. It was possible to verify that those species co-occurred with amentoflavone in the same study and investigate whether a plant species could produce amentoflavone (Supplementary Table S1).

**Figure 2.**
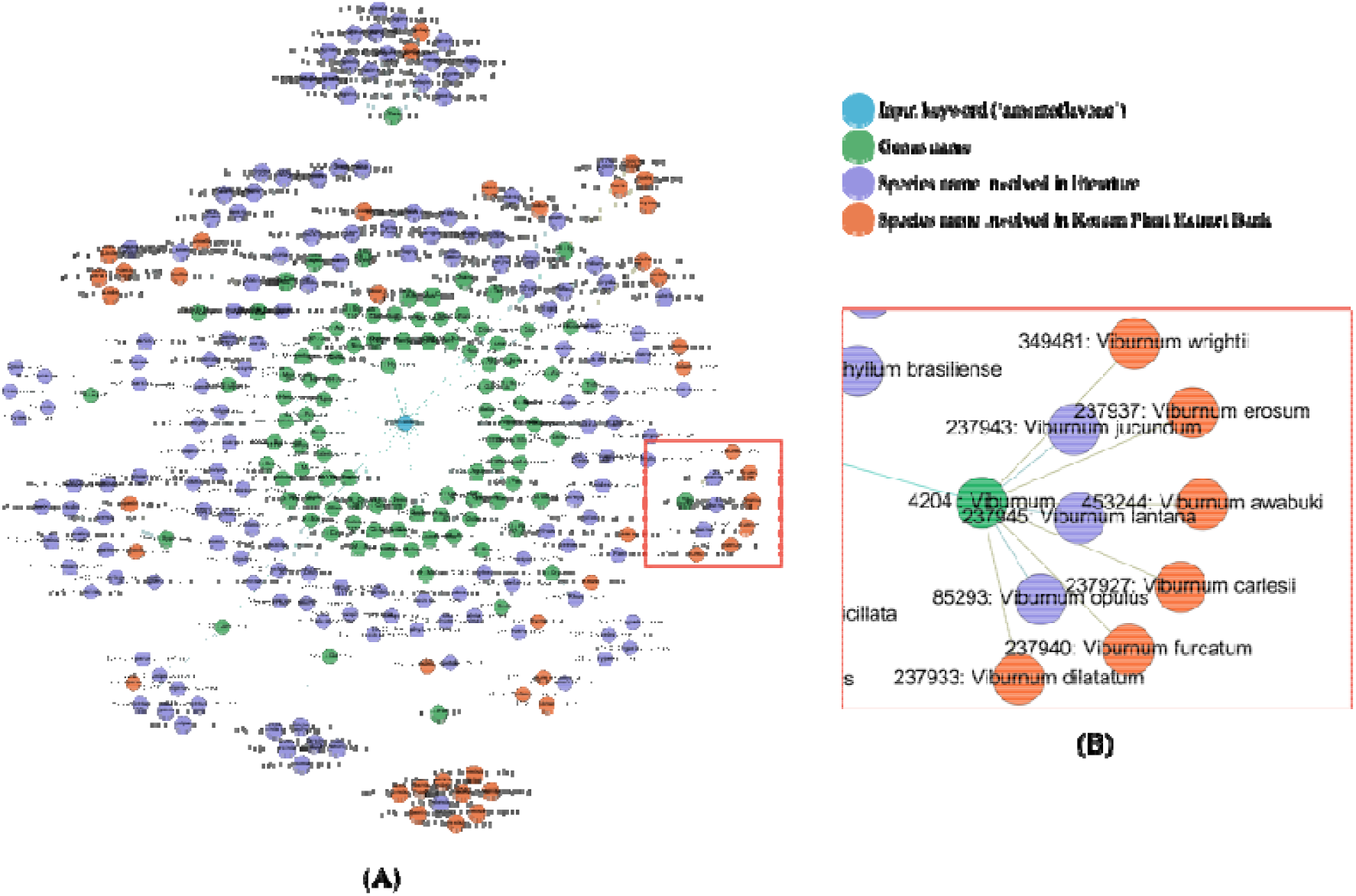
(A) Network obtained by entering ‘amentoflavone’ as input keyword in ChexMix. The unique identifiers (TaxID, purple nodes) for species co-existing with the input keyword in the literature are linked to their own taxonomic higher rank (genus, green color). Orange nodes represent species names that only existed in the list of Korean medicinal plants of the KPEB and are linked to the nodes for genus to which each species belongs. (B) Detailed subnetwork under the *Viburnum* genus. Each node was displayed as ‘ID: name’ for TaxID and genus or species name.

To avoid duplicated studies and find novel bioactive sources, the analysis was focused on the allied species belonging to the *Viburnum* genus, retrieving 19 samples of different parts of eight species native to Korea that were not previously studied on amentoflavone-related topics (Fig. 3, Supplementary Table S2). Next, the existence of amentoflavone was evaluated in samples of these plants and quantified by HPLC. The presence of amentoflavone was confirmed by its isotopic peak at 537.4 m/z [M + H]^−^ detected by liquid chromatography–mass spectrometry. Among them, the leaves of *V. erosum* contained the highest amount of amentoflavone (7.39 mg/g) compared with *Selaginella tamariscina*, which is the representative natural ingredient for anti-wrinkle effect and the major source of amentoflavone in the cosmetic industry^19^. Overall, the summarization of hierarchical bioentities information using ChexMix is expected to help inspect massive sparse bioentities in databases in future investigations.

**Figure 3.**
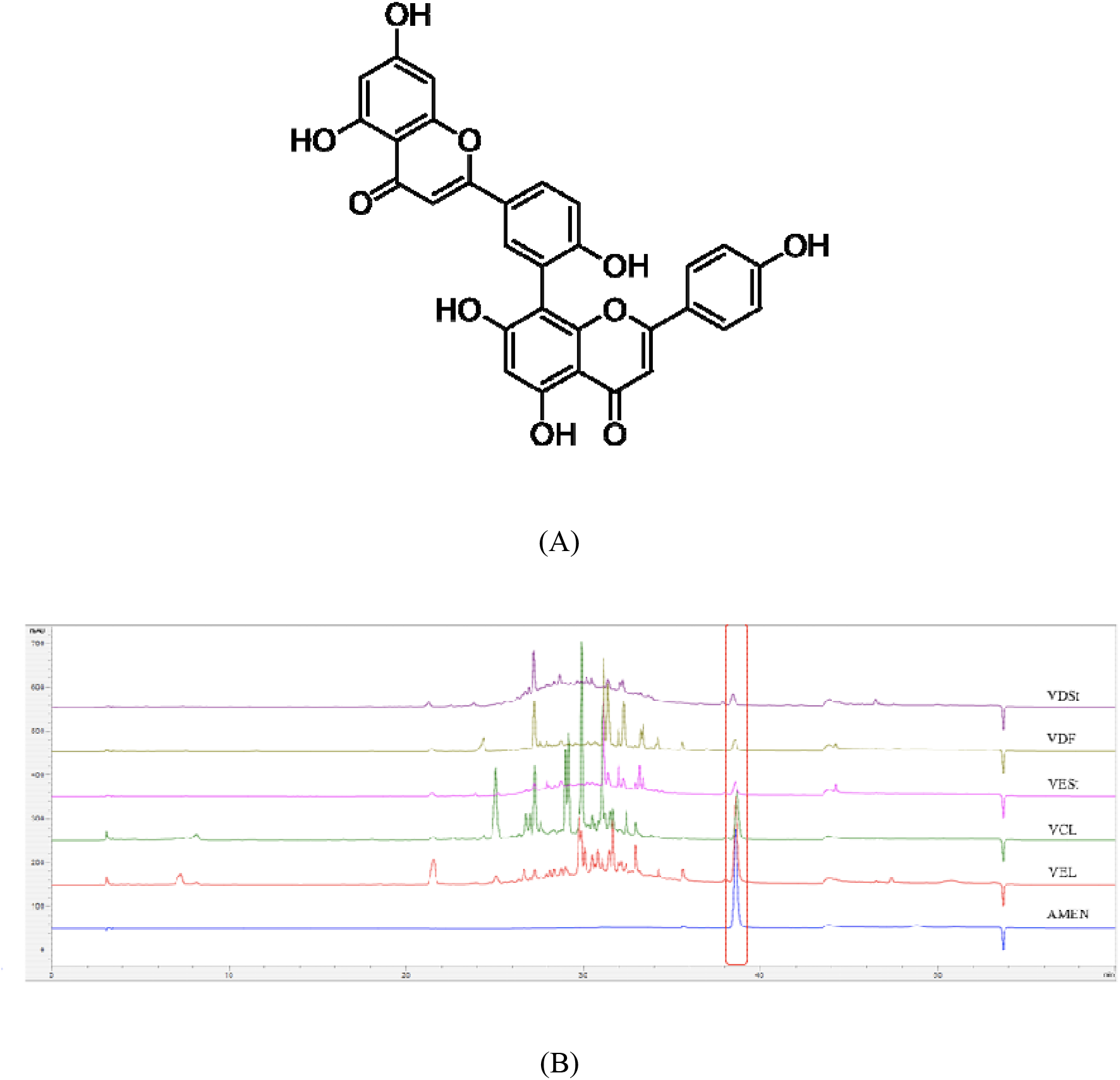
(A) Chemical structure of amentoflavone. (B) Chromatograms of the five samples with the highest amentoflavone content determined as described in the ‘Methods’ section. AMEN, amentoflavone; VCL, leaves of *Viburnum carlesii;* VDF, fruits of *V. furcatum*; VDSt, leaves of *Viburnum dilatatum;* VEL, leaves of *V. erosum*; VESt, stems of *V. erosum*;.

Additionally, ChexMix can also integrate the results from multi-keywords. The MeSH identifiers for bioentities co-occurring with the keywords of interests could be used for connecting the results by two different queries (Fig. 4). For instance, two species names, *Taxus cuspidata* and *Podophyllum peltatum*, were queried by ChexMix and generated two small networks consisting of bioentities with MeSH identifiers extracted from PubTator. It was possible to inspect the co-occurred bioentities among the MeSH identifiers in the integrated network. The network of each species showed different MeSH identifier profiles and MeSH identifiers related to ‘cancer’, in particular ‘ovarian neoplasms’, co-occurred. This agrees with the fact that paclitaxel of *T. cuspidata* and podophyllotoxin of *P. peltatum* are well-known potent anticancer drugs for ovarian cancer^20–22^.

**Figure 4.**
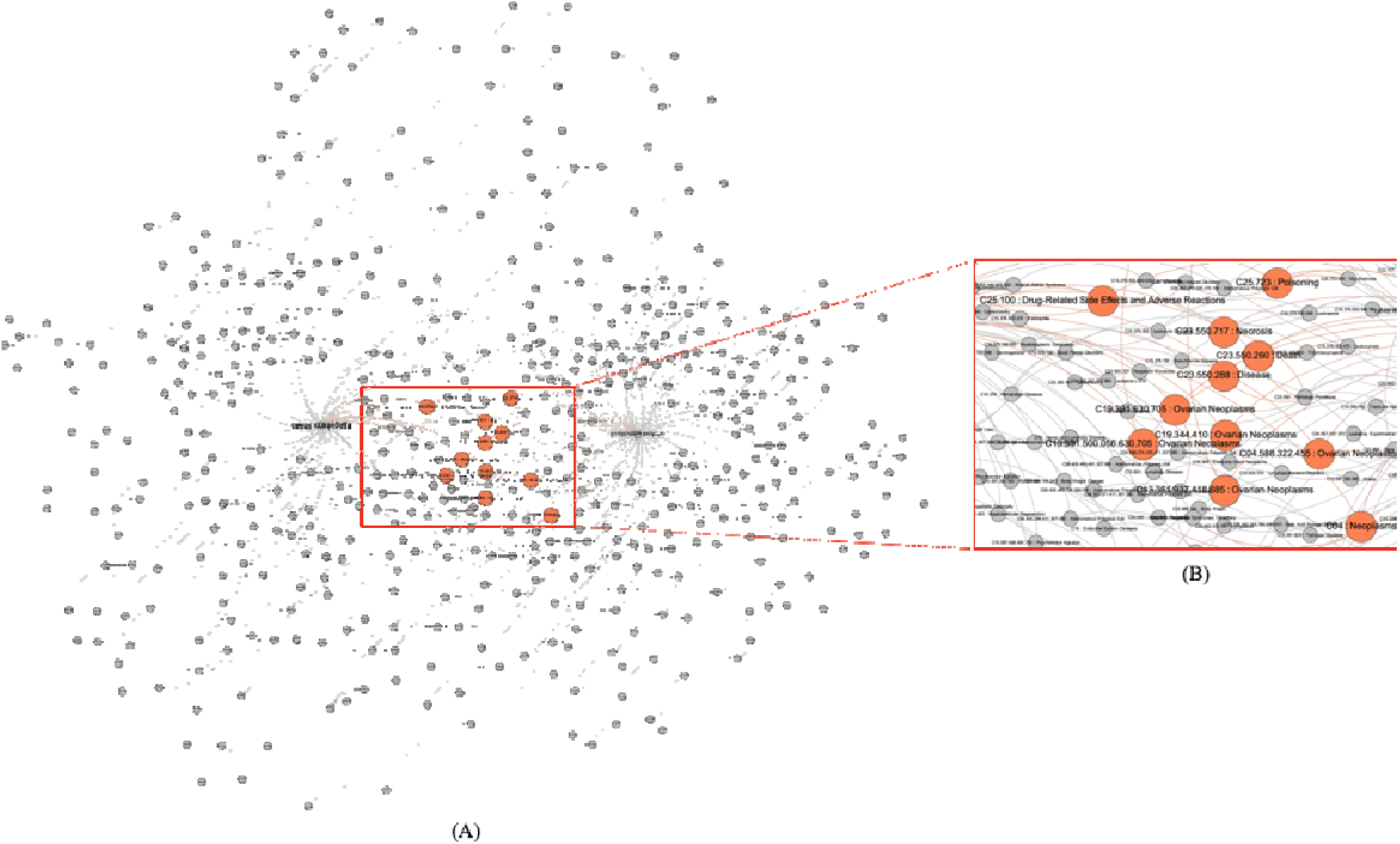
(A) Acquired network using *‘taxus cuspidata’* and *‘Podophyllum peltatum’* as input keywords in ChexMix. MeSH terms co-occurring in the literature with the input keywords were reorganized according to the hierarchy rules of the MeSH Tree Structures in the MeSH browser (https://meshb-prev.nlm.nih.gov/treeView). The nodes of the co-occurred bioentities in both keywords are colored in orange. (B) Details of the subnetwork of the co-occurred bioentities in both keywords. Each node displays as ‘Tree Number: MeSH Heading’ for MeSH identifiers and a MeSH term.

Here, a usage scenario of ChexMix to alleviate the complex task of compiling large data by narrowing down the scope of bioentities or grouping similar bioentities using the hierarchical relationships was described. Firstly, to obtain the appearance counts of bioentities in literature queried by keywords of interest, ChexMix collects PubMed and PMC literature followed by fetching annotations within that data from PTC and converting them into unique identifiers according the respective bioentity class. ChexMix allows Boolean operators (‘AND’, ‘OR’, ‘NOT’), double quotes for phrases, and asterisk for truncated terms for PubMed literature search. Each bioentity extracted from ChexMix is classified within more general categories of bioentity and arranged in a hierarchical structure.

When single or multi keywords of interest are entered in ChexMix, bioentities in all citations that have keywords are retrieved and automatically mapped into unique identifiers. The search results indicate the co-occurrence of bioentities in the available literature, allowing to link them and yielding the co-occurrence network. ChexMix makes the process straightforward by managing data access from multiple sources and providing functions to manipulate the network data structure.

The analysis is mainly focused on taxonomy terms to inspect the species that biologically affect physiological disorders or diseases within the network. Each taxonomy name in the search results is listed in a hierarchical form. Trivial bioentities are located on the higher ranks of the list. Other near species within the obtained taxonomic tree are expected to have similar biological effects, representing potential alternative biomedical options. ChexMix can also generate the connections between taxonomic terms and MeSH identifiers, which are located under ‘Diseases [C]’ and ‘Chemicals and Drugs [D]’, in the same literature. MeSH identifiers co-occurring with a taxonomic term in the literature are expected to have a close relationship.

In Fig. 4, the intersection set of MeSH terms co-occurring with each taxonomy keyword is highlighted on the whole network resulting from the union set of two networks. Networks generated from a single keyword in ChexMix can be simply reprocessed by the combination of set operations, such as union, difference, and intersection with other networks. The reorganization of complex networks from single or multi keywords provides new insights or clues for bioentities in PubMed, the biggest biomedical database.

## Methods

### Data Processing

ChexMix currently obtains biomedical data from multiple databases using their web application programming interface (APIs) or bulk data files. For example, Entrez API allows to query Entrez databases, such as PubMed and PubMed Central (PMC), using combinations of keywords. PTC provides a web API to fetch annotations of biomedical concepts, such as taxonomy and MeSH identifiers, in a publication. ChexMix also manages to download and parse bulk data files from biomedical databases^16^. For example, ChexMix loads the data from PTC, including NCBI taxonomy and MeSH that inherently have relationships between entities therein, and transforms it into internal network data structures. ChexMix also grants the possibility to construct, manipulate, and simplify the network data structures.

### Bioentities extraction and visualization

The keywords of interest can be input as single words or phrases. The results are output in hierarchical tree format according to their own taxonomic or hierarchically-organized rules for each type of bioentity. In the case of taxonomy information, species names in the literature are encoded into unique identifiers, TaxID, and hierarchically re-organized in the classification rules of NCBI taxonomy. In the present study, hierarchical results were applied to discover relevant species with lower taxonomic ranks (family and genus levels) using the list of the Korean medicinal plants of the Korea Plant Extract Bank (KPEB). The results were visualized in the network format using the Gephi software (ver. 0.9.2).

### Sample preparation

To prove the usefulness of ChexMix, 19 *Viburnum* samples, including *V. erosum*, *V. carlesii*, *V. dilatatum, V. wrightii, V. sargentii, V. opulus, V. furcatum*, and *V. awabuki*, were collected from the Medicinal Herb Garden of the College of Pharmacy, Seoul National University, Korea, or purchased from the KPEB of the Korea Research Institute of Bioscience and Biotechnology, Korea. Each powdered sample (1 g) was extracted using 80% methanol for 1 h using an ultrasonic apparatus and the filtrate was dried using a vacuum rotary evaporator. Next, the samples were dissolved in 100% methanol at a concentration of 10 mg/mL and filtered through a 0.45 μm polytetrafluoroethylene membrane before analysis.

### High-performance liquid chromatography (HPLC) analysis

The samples were analyzed on a 1260 quaternary pump, an autosampler, and a multiple wavelength detector (Agilent Technologies, Santa Clara, CA, USA). Chromatographic separation was performed using a Hector□M C18 column (250 × 4.6 mm I.D.; 5 μm, RSTech, Daejeon, Korea). The ultraviolet detector was set at a wavelength of 260 nm. The mobile phase was a gradient solvent system consisting of solvent A (0.1% formic acid in water) and solvent B (MeOH) as follows: isocratic 95% A (0–10 min), linear gradient 95–80–30% A (10–20–30 min) and isocratic 30% A (30–40 min). The flow rate was 1.0 mL/min, and aliquots of 10 μL were injected using an autosampler.

## Code Availability

The in-house codes used to extract the bioentities and classify them, as well as to perform the data visualization, along with some examples, are publicly available at the github repository: https://github.com/bionsight/chexmix.

## Acknowledgements

This work was supported by the Basic Science Research Program through the National Research Foundation of Korea (NRF) grant funded by the Korean government (MEST) (NRF- 2021R1C1C1011857).

## Author Contributions

HeejungY. and Hojin.Y. conceived and coordinated the project. J.P., J.L. and H.S.J. collected *Viburnum* samples and analyzed the content of amentoflavone. B.P., Heejung Y. and Hojin Y. wrote the in-house python codes. B.P, N.L., Hojin Y. and Heejung Y. contributed to discussions, and Heejung Y. and Hojin Y. wrote the manuscript.

---

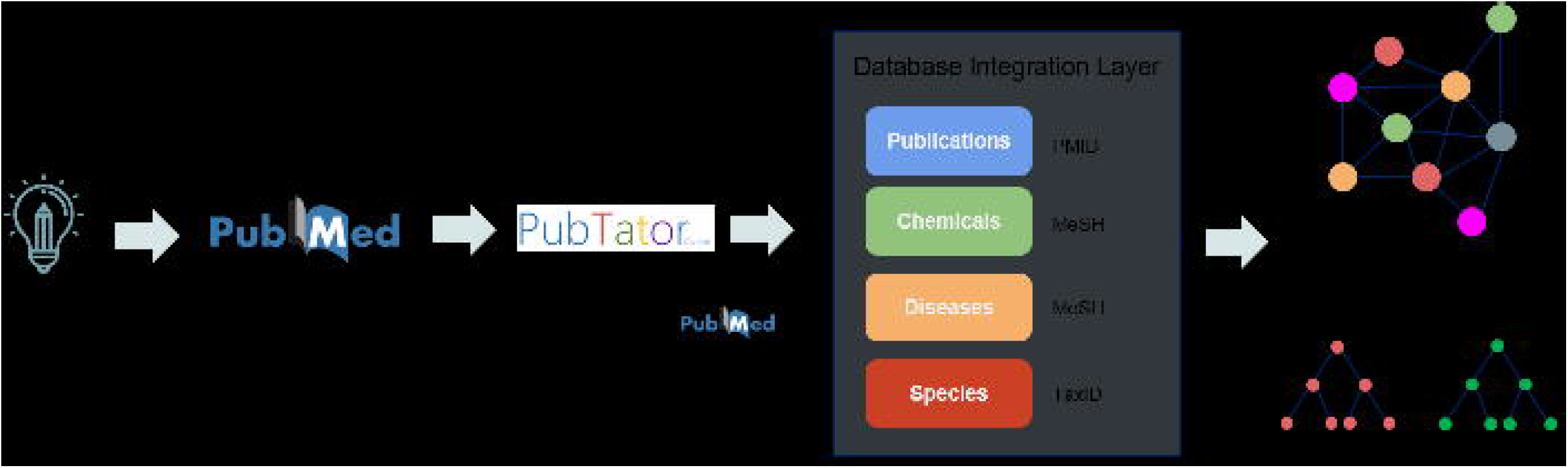

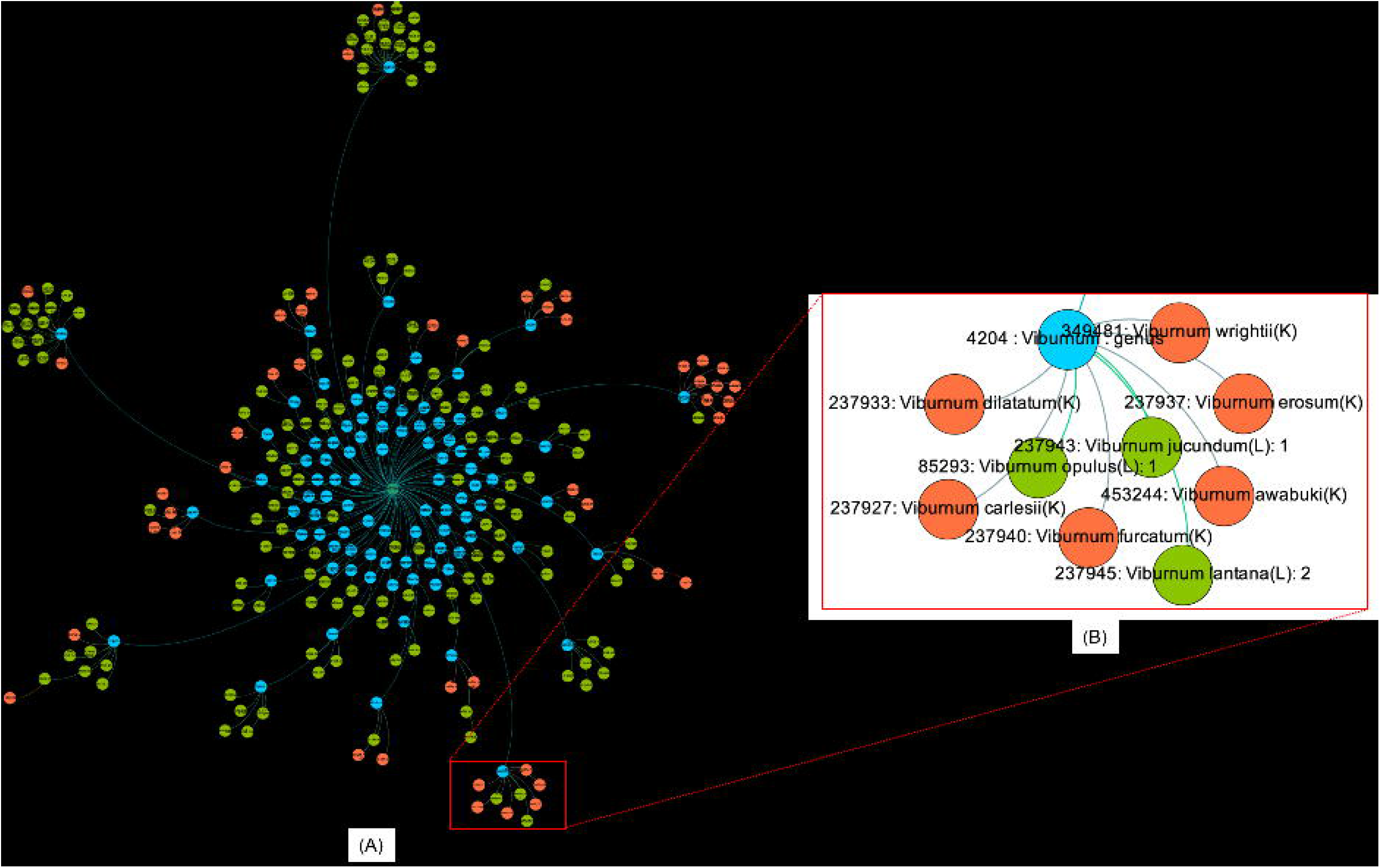

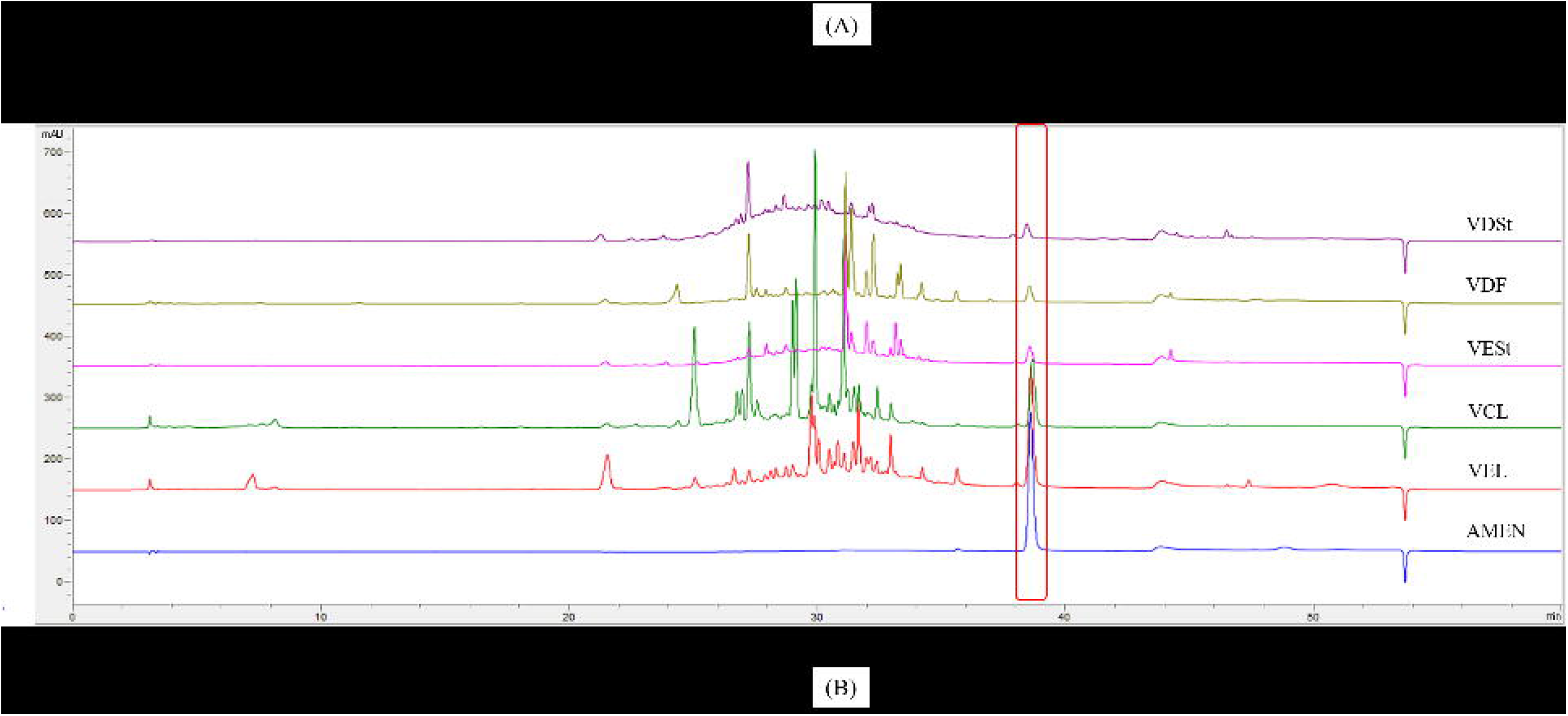

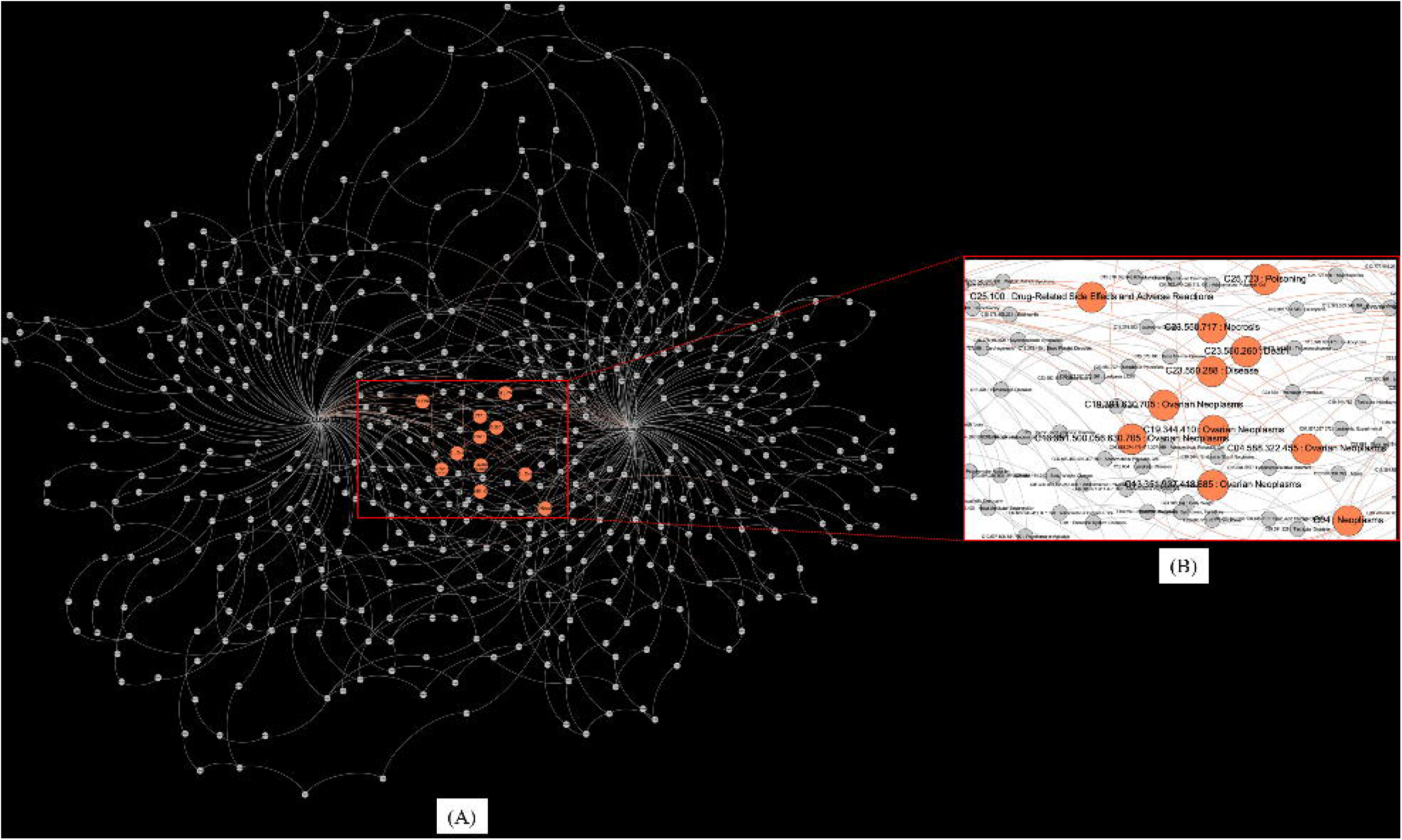

## References

1. Mendez, D. et al. ChEMBL: towards direct deposition of bioassay data. Nucleic Acids Res. 47, D930–D940 (2019).

2. Gilson, M. K. et al. BindingDB in 2015: A public database for medicinal chemistry, computational chemistry and systems pharmacology. Nucleic Acids Res. 44, D1045–D1053 (2016).

3. Wassermann, A. M. & Bajorath, J. BindingDB and ChEMBL: online compound databases for drug discovery. Expert Opin. Drug Discov. 6, 683–687 (2011).

4. Wishart, D. S. et al. DrugBank 5.0: a major update to the DrugBank database for 2018. Nucleic Acids Res. 46, D1074–D1082 (2018).

5. Davis, A. P. et al. The Comparative Toxicogenomics Database: update 2019. Nucleic Acids Res. 47, D948–D954 (2019).

6. Wilson, S. et al. Automated literature mining and hypothesis generation through a network of Medical Subject Headings. (2018) doi:10.1101/403667.

7. Himmelstein, D. S. et al. Systematic integration of biomedical knowledge prioritizes drugs for repurposing. eLife 6, e26726 (2017).

8. Wei, C.-H., Kao, H.-Y. & Lu, Z. PubTator: a web-based text mining tool for assisting biocuration. Nucleic Acids Res. 41, W518–W522 (2013).

9. Wei, C.-H., Allot, A., Leaman, R. & Lu, Z. PubTator central: automated concept annotation for biomedical full text articles. Nucleic Acids Res. 47, W587–W593 (2019).

10. Federhen, S. The NCBI Taxonomy database. Nucleic Acids Res. 40, D136–D143 (2012).

11. Ertl, P. An algorithm to identify functional groups in organic molecules. J. Cheminformatics 9, (2017).

12. Ertl, P. & Schuhmann, T. Cheminformatics Analysis of Natural Product Scaffolds: Comparison of Scaffolds Produced by Animals, Plants, Fungi and Bacteria. bioRxiv 2020.01.28.922955 (2020) doi:10.1101/2020.01.28.922955.

13. Djoumbou Feunang, Y. et al. ClassyFire: automated chemical classification with a comprehensive, computable taxonomy. J. Cheminformatics 8, 61 (2016).

14. Uhlen, M. et al. Tissue-based map of the human proteome. Science 347, 1260419–1260419 (2015).

15. Uhlen, M. et al. A pathology atlas of the human cancer transcriptome. Science 357, eaan2507 (2017).

16. Swainston, N. et al. libChEBI: an API for accessing the ChEBI database. J. Cheminformatics 8, 11 (2016).

17. Yu, S. et al. A Review on the Phytochemistry, Pharmacology, and Pharmacokinetics of Amentoflavone, a Naturally-Occurring Biflavonoid. Molecules 22, 299 (2017).

18. Park, N.-H., Lee, C.-W., Bae, J. & Na, Y. J. Protective effects of amentoflavone on Lamin A-dependent UVB-induced nuclear aberration in normal human fibroblasts. Bioorg. Med. Chem. Lett. 21, 6482–6484 (2011).

19. Yuan, C. Simultaneous determination of selaginellins and biflavones in Selaginella tamariscina and S. pulvinata by HPLC. China J. Chin. Mater. Medica (2012) doi:10.4268/cjcmm20120918.

20. Baird, R. D., Tan, D. S. P. & Kaye, S. B. Weekly paclitaxel in the treatment of recurrent ovarian cancer. Nat. Rev. Clin. Oncol. 7, 575–582 (2010).

21. Zhao, W. et al. Challenges and potential for improving the druggability of podophyllotoxin-derived drugs in cancer chemotherapy. Nat. Prod. Rep. 10.1039.D0NP00041H (2021) doi:10.1039/D0NP00041H.

22. Mukherjee, A., Basu, S., Sarkar, N. & Ghosh, A. Advances in Cancer Therapy with Plant Based Natural Products. Curr. Med. Chem. 8, 1467–1486 (2001).

