## Supplemental for "ChexMix: A Literature Content Extraction Tool for Bioentities": Supplementary_Tables.docx

Table S1. The results for an input keyword, ‘amentoflavone’, in ChexMix^TM^

| Taxonomy ID | Species | pmid |
| --- | --- | --- |
| 4113 | *Solanum tuberosum* | 30612057 |
| 3750 | *Malus domestica* | 30236673 |
| 3311 | *Ginkgo biloba* | 32144952, 31913733, 31526502, 31446241, 30995808, 30693991, 19356077, 17225453, 16557462, 16432273, 16084098, 15285849, 12824018, 12622229, 11853165, 11842336, 9834158, 3212094 |
| 169191 | *Schinus terebinthifolia* | 32466911, 24881808 |
| 3882 | *Onobrychis viciifolia* | 21930278 |
| 261999 | *Camellia sinensis* | 28554631 |
| 4270 | *Byrsonima crassifolia* | 18361746, 17459621 |
| 137178 | *Selaginella tamariscina* | 32343974, 32180727, 31699478, 31549342, 30507085, 29295657, 28942285, 28603761, 28286104, 28185818, 27106512, 25422557, 25240932, 24358788, 24280521, 23970815, 23727809, 23160679, 22803371, 22210020, 21710597, 21663495, 19652385, 18591814, 18479689, 18029185, 17917274, 17268085, 17024847, 16289413, 15829434, 15326548, 8792657 |
| 2291027 | *Salvia incertae sedis* | 17923139 |
| 179351 | *Dendrobium capillipes* | 27353868 |
| 85293 | *Viburnum opulus* | 21186982 |
| 237943 | *Viburnum jucundum* | 11582553 |
| 237945 | *Viburnum lantana* | 21186982, 3002388 |
| 190902 | *Mentha aquatica* | 18775771 |

Table S2. The contents of amentoflavone in 8 Viburnum species

| Species | Sample Name | Parts | Contents (mg/g) | Collection location |
| --- | --- | --- | --- | --- |
| *V. erosum* | VEL | leaves | 7.39 | SNU |
|  | VESt | stems | 1.40 | KRIBB |
|  | VEBc | branches | n.d | KRIBB |
| *V. carlesii* | VCL | leaves | 4.56 | SNU |
|  | VCSt | stems | 0.47 | KRIBB |
| *V. dilatatum* | VDSt | stems | 1.25 | KRIBB |
|  | VDL | leaves | 0.78 | KRIBB |
|  | VDF | flowers | 1.36 | KRIBB |
|  | VDBc | branches | n.d | KRIBB |
| *V. wrightii* | VWW | whole | 0.55 | KRIBB |
| *V. sargentii* | VSF | fruits | n.d | KRIBB |
|  | VSBc | branches | n.d | KRIBB |
|  | VSL | leaves | n.d | KRIBB |
|  | VSSd | seeds | n.d | KRIBB |
|  | VSSt | stems | n.d | KRIBB |
| *V. opulus* | VOL | leaves | 0.70 | SNU |
|  | VOBc | branches | n.d | SNU |
| *V. furcatum* | VFL | leaves | n.d | KRIBB |
|  | VFBk | barks | n.d | KRIBB |
|  | VFS | stems | n.d | KRIBB |
| *V. awabuki* | VAL | leaves | n.d | SNU |
|  | VABc | branches | n.d | SNU |
